# Oncogenic PTPN11/SHP2 drives immune escape in juvenile myelomonocytic leukemia (JMML) through activation of ectonucleotidase/adenosine signaling

**DOI:** 10.1101/2025.10.01.679664

**Authors:** Jovana Rajak, Ke Meng, Anna Lena Stippel, Jun Wang, Naile Koleci, Sheila Bohler, Alexandra Emilia Schlaak, Hui Xiao, Lukas M. Braun, Riccardo Masetti, Robert Zeiser, Christian Flotho, Daniel B. Lipka, Charlotte M. Niemeyer, Luciana Hannibal, Bertram Bengsch, Miriam Erlacher

## Abstract

Juvenile myelomonocytic leukemia (JMML) is a myelodysplastic/myeloproliferative neoplasm of early childhood driven by RAS pathway mutations. Allogeneic hematopoietic stem cell transplantation (HSCT) is the therapy of choice for most patients. However, relapse rate is high, in patients with adverse features, frequently noted in *PTPN11*-mutated JMML, or in patients without evidence of graft-versus-host disease (GvHD). Here we set out to understand the mechanisms associated with oncogenic *PTPN11* immune escape. Analyzing primary *PTPN11*-mutated JMML samples and *MxCre;Ptpn11^D61Y/+^* mice, we observed elevated expression of immune checkpoint molecules, including ectonucleotidases CD39 and CD73 - key mediators of the adenosine pathway - on monocytic and granulocytic leukemic cells. Stimulation with GM-CSF, a central mediator of JMML pathogenesis, induced ectonucleotidases expression on granulocytes and monocytes. In contrast, MEK inhibition downstream of *Ptpn11^D61Y/^*^+^ reduced ectonucleotidases expression. Functionally, *Ptpn11^D61Y/^*^+^-mutated myeloid cells suppressed activation and proliferation of wild-type (WT) T lymphocytes, an effect recapitulated by adenosine and reversed by pharmacological CD39 inhibition with POM-1. *In vivo*, POM-1 treatment of *MxCre*;*Ptpn11^D61Y/+^*mice presenting with myeloproliferation reduced spleen size and partially restored immune responsiveness. Moreover, POM-1 induced apoptosis in murine *Ptpn11^D61Y/+^* myeloid cells, highlighting a dual therapeutic benefit of CD39 inhibition in JMML. Together, these findings suggest that targeting the adenosine pathway may represent an immunomodulatory approach to enhance T cell-mediated control of JMML, particularly in the context of HSCT and relapse prevention.

## INTRODUCTION

Juvenile myelomonocytic leukemia (JMML) is a rare myelodysplastic/myeloproliferative neoplasm of early childhood (1, 2). JMML is characterized by an abnormal cell proliferation of the granulocytic-monocytic lineage, causing organ infiltration with splenomegaly as the most prominent clinical sign (1). The key oncogenic driver in the pathogenesis of the disease is the constitutive activation of the RAS/MAPK signaling pathway caused by mutations in *PTPN11*, *KRAS*, *NRAS*, *NF1* and *CBL* (3, 4). For most patients, allogeneic hematopoietic stem cell transplantation (HSCT) is the treatment of choice. However, in up to 35% of patients, disease eradication fails and JMML relapses (5, 6). JMML recurrence is inversely correlated with the presence and severity of chronic graft-versus-host disease (cGVHD), indicating that a graft-versus-leukemia (GvL) effect plays a major role in eradicating the disease (6–8). Accordingly, rapid tapering of immunosuppressive therapy to enhance GvL is recommended in JMML patients with high-risk features (9, 10).

Immune escape mechanisms, i.e. strategies to evade GvL, play a critical role in leukemia recurrence after HSCT (11). While immune evasion has been extensively studied in acute myeloid leukemia (AML) and is increasingly recognized in acute lymphoblastic leukemia (ALL) (12–15), its role in JMML remains largely unexplored.

*PTPN11*, which encodes the oncogenic protein tyrosine phosphatase SHP2, is mutated in up to 40% of JMML patients (4). Many patients with JMML driven by somatic *PTPN11* mutations are diagnosed at ≥ 2 years of age, with elevated fetal hemoglobin levels ≥ 15%, and a hypermethylated DNA profile, all of which are known to be associated with a high-risk of disease recurrence after HSCT (16–18). In contrast to somatic mutations, germline mutations in *PTPN11* cause Noonan syndrome (NS) and may lead to a transient myeloproliferative disorder (MPD) early in life (3). Beyond driving hematopoietic stem and progenitor cell (HSPC) proliferation and differentiation (19), oncogenic SHP2 signaling has also been implicated in promoting tumor evasion (20–22). Yet, the role of SHP2 in regulating immune checkpoint pathways and their functional relevance in JMML has to be elucidated.

Using primary JMML cells and a mouse model reminiscent of human JMML, we identify elevated expression of multiple immune checkpoint molecules. Amongst those, the ectonucleotidases CD39 and CD73 were directly regulated by the oncogenic SHP2 signaling. Murine *Ptpn11^D61Y/+^*mutated myeloid cells severely inhibited proliferation and activation of T cells, a phenomenon that could be reversed by CD39 inhibition. In addition, CD39 blockade induced cell death in murine *Ptpn11^D61Y/+^* myeloid cells, indicating dual function of this ectonucleotidase in JMML. Our findings suggest that therapeutic inhibition of CD39 may hold promise as a targeted treatment strategy.

## METHODS

### Primary human cells

Patients had been included in studies EWOG-MDS 98 (#NCT00047268) and EWOG-MDS 2006 (#NCT00662090) of the European Working Group of MDS in Childhood (EWOG-MDS). The studies had been approved by the institutional ethics committees of the respective institutions. Written informed consent was obtained from patients’ guardians. Clinical details for patients studied are listed in Suppl. Table 1. JMML spleen cells were obtained from splecetomized spleens. Spleen tissue was mechanically dissociated to obtain single-cell suspensions. Mononuclear cells (MNCs) were subsequently isolated by Ficoll-based density gradient centrifugation and cryopreserved for further analyses.

### Mass Cytometry

#### Preparation of single-cell suspensions

Splenocytes from 10 JMML patients with somatic *PTPN11* mutation, 2 patients with Noonan syndrome and transient MPD, and 4 controls were thawed in RPMI-1640 medium supplemented with 10% FCS, 1% P/S, 10nM HEPES and 2 ng/mL DNase I, centrifuged at 300g for 5 minutes. After a 10-minute wash in fresh RPMI3+, cell pellets were resuspended in the same medium for staining.

#### Staining procedure

A 46-marker panel was used (Suppl. Table 2). Staining were performed as described (23). Briefly, pellets were stained with a live/dead buffer at room temperature (RT) for 5 minutes, washed twice, and barcoded using three unique combinations of 104Pd, 105Pd, 106Pd, 108Pd, 110Pd, and 198Pt at 4°C for 30 minutes. After two washes, samples were pooled, incubated with extracellular antibody cocktail at RT for 30 minutes, washed twice, fixed, permeabilized, and stained intracellularly. After two final washes, cells were incubated overnight at 4°C in 4% PFA solution containing 125nM 191/193 Iridium. Samples were frozen until acquisition on a Helios CyTOF system (Fluidigm).

#### Data preprocessing and analysis

Bead-based normalization corrected signal drift over time. Debarcoding was performed in FlowJo (v10.7.1) according to the barcoding schema. FCS files were processed in Omiq.ai platform. Sequential gating removed aggregates, debris, and doublets, identifying singlet CD45-positive cells. Manual gating compared cell composition between JMML *PTPN11* mutant and control samples. Unsupervised analyses assessed immune composition, checkpoints, and regulatory molecules in JMML. Additional information on sample processing, sequencing, and computational analyses is provided in the Supplementary Methods.

### Mice

*Ptpn11^D61Y/+^* mice were generated by Benjamin Neel and bred with *MxCre* to obtain *MxCre;Ptpn11^D61Y/+^* mice (24). To induce *Ptpn11^D61Y^* expression, mice received intraperitoneal (i.p.) injections of 300 µg of polyinosinic-polycytidylic acid (poly(I:C); Sigma-Aldrich) every other day for three doses. *MxCre* and *Ptpn11^D61Y/+^*mice were maintained on a C57BL/6N background and genotyped as described (Suppl. Table 3) (22). B6.SJL-Ptprca Pepcb/BoyJ mice served as splenic T-cells donors for transplantation experiments. All animals were housed under specific-pathogen-free conditions. Experiments were conducted in accordance with Animal Welfare Regulations.

### *In vivo* treatment with the CD39 inhibitor POM-1

*MxCre* and *MxCre;Ptpn11^D61Y/+^* mice were treated with poly(I:C). CD39 inhibition was performed with POM-1 administered every other day for 14 days (7 doses, 25/mg/kg/day). Mice were sacrificed one day after the final treatment, and BM and spleen were collected for analysis of hematopoietic cell compartments by flow cytometry. Detailed protocol is available in the Supplementary Methods.

### Organ analysis and cell staining

BM was collected by flushing femora, tibiae, and pelvis with PBS containing 5 IU/mL penicillin, 5 µg/mL streptomycin (P/S), and 10% fetal-calf serum (FCS) using a 27G needle. Single-cell suspensions from the spleens were prepared by mechanical dissociation through a 70 μm and 40 μm strainer due to high cellularity strainer. Blood, BM, and spleen samples were subjected to red blood cell lysis (150 mM NH₄Cl, 10 mM NaHCO₃, 1 mM EDTA) for 10 minutes (min) at RT. Cell counts were determined with a Neubauer chamber; BM counts were based on two femora. Cells were stained with surface antibodies (1:100) in the presence of FcR block (1:100), and dead cells were excluded using a viability dye (1:800). Data were acquired on an LSR Fortessa and analyzed using FlowJo (BD Biosciences). Gating strategy is shown in Supplementary Figure 2. and staining was performed using antibodies provided in Supplementary Table 4.

### Quantification of adenosine, AMP, ADP and ATP in murine plasma

Mouse plasma samples were collected in EDTA-coated tubes and stored at –80 °C. Stock solutions of adenosine, AMP, ADP, and ATP were prepared in LC-MS-grade water and stored at –80 °C. Calibration curves (0–50 µM) were generated by serial dilution with water. For each 20 µL sample, 20 µL of 15N5-ATP (0.5 µM) and 20 µL LC-MS-grade water were added, vortexed at RT, then extracted with 100 µL methanol, vortexed, and centrifuged (10,000 × g, 10 min, RT). 110 µL of supernatant was transferred to HPLC vials. Separation was performed on an X-Terra C18 column using a gradient of solvent A (4 mM ammonium formate in water) and solvent B (4 mM ammonium formate in 90% acetonitrile). Data were acquired and quantified using Analyst® software v1.7.2 (AB Sciex, 2022). Detailed protocol is available in the Supplementary Methods.

### Myeloid cells isolation and culture

CD11b⁺ myeloid cells were isolated from mouse spleens using magnet-activated cell sorting (MACS) with the CD11b MicroBeads kit (Miltenyi Biotec). For further separation of monocytic and granulocytic subsets, cells were stained with antibodies against Ly6C and Ly6G together with a viability dye to exclude dead cells and sort accordingly. Sorted cells (1×10⁵ per well) were seeded into 96-well U-bottom plates and cultured for either 24 or 96 hours. Detailed cell culture protocol is available in the Supplementary Methods.

### *In vitro* ATP hydrolysis–based measurement of ATPase activity

ATPase activity was assessed by measuring extracellular ATP hydrolysis. Spleen cells were cultured in complete medium with or without POM-1 (50 μM) for 16 hours. After washing, the cells were incubated in fresh medium again with or without POM-1 (50 μM) in the presence of 10 μM ATP for 30 minutes at 37°C. Extracellular ATP levels were quantified using the ATPlite Luminescence ATP Detection Assay System (PerkinElmer).

### *In vitro* T cell proliferation and activation assays

αCD3/CD28-mediated T cell stimulation was performed using splenic T cells isolated from WT mice via the Pan T Cell Isolation Kit (Miltenyi Biotec). Purified 10⁵ T cells were co-cultured with Dynabeads™ Mouse T-Activator CD3/CD28 either alone or in 1:1 ratio with monocytes or granulocytes. T cell proliferation was assessed using CFSE, and cells were harvested on day 4 for analysis. T cell proliferation and activation were analyzed by flow cytometry using CFSE and CD44/CD62L staining, respectively. Where indicated, cultures were exposed to POM-1, APCP, anti-PD-1, or anti-VISTA; full compound and antibody information is listed in the Supplementary Methods.

### Statistics

Statistical analyses of mass cytometry data were performed using R (v4.4.2) and GraphPad Prism 10. Data are presented as mean ± SEM, and significance was defined as *p* < 0.05. A full description of statistical tests is available in the Supplementary Methods.

## RESULTS

### CyTOF-based immune profiling of JMML patients harboring a *PTPN11* mutation

We investigated the immunoregulatory landscape of human JMML using comprehensive immunoprofiling of diseased spleens. A 46-plex mass cytometry panel was established for the detailed profiling of human adaptive and innate immune subsets, JMML cells, differentiation markers and immune checkpoints (Supplementary Table 2). Spleens from 10 JMML patients harboring a *PTPN11* mutation and 2 Noonan syndrome patients with germline *PTPN11* mutation were analyzed. As controls, spleens from 4 patients with non-malignant disease (i.e. patients with hemoglobinopathies or hereditary spherocytosis) were included. Samples were acquired using mass cytometry (Figure 1A). Data preprocessing strategies and manual gating strategies are provided in Supplementary Figure 1A-B.

**Figure 1.**
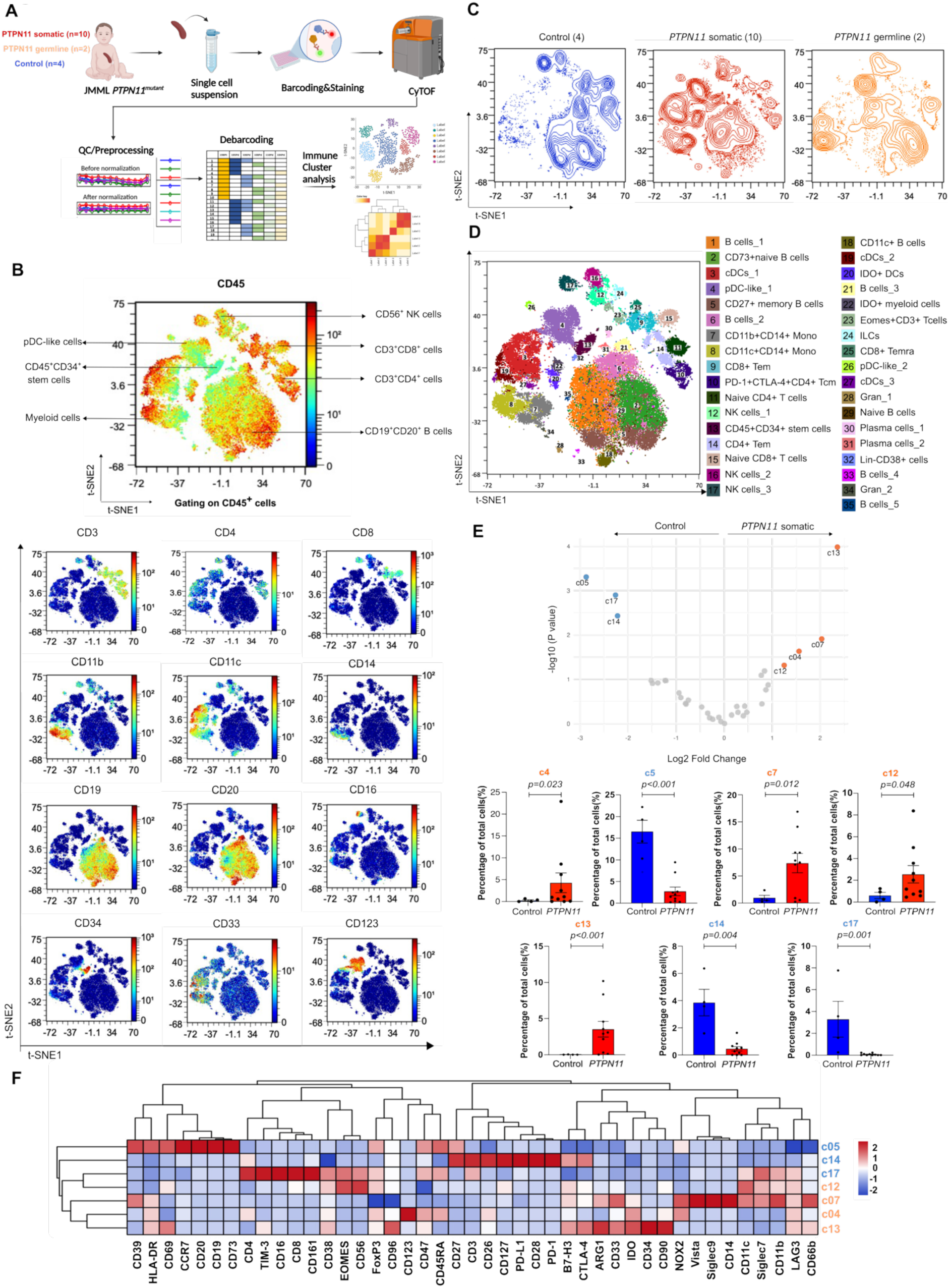
Unsupervised analysis reveals detailed distinct pattern of immune dysregulation in JMML splenocytes. **(A)** Workflow of CyTOF staining and data analysis. **(B)** Coloured-continuous scatterplot showing lineage expression markers by a dimensionality reduction t-SNE map on all samples. Expression markers are shown below. **(C)** Contour plot t-SNE showing immune distribution of experimental groups: control (n=4), *PTPN11* somatic (n=10) and *PTPN11* germline (n=2). **(D)** PARC clustering on all samples resulting in 35 clusters. (**E)** Vocano plot and box plots of significant clusters. Volcano plot illustrating the differential expression analysis between control group and JMML *PTPN11*-mutant group. Each point represents a cluster, with the y-axis showing the fold change (log2 scale) and the x-axis representing the statistical significance (-log10 p-value). Significant clusters are highlighted in orange (upregulated) or blue (downregulated) above the cut-off of fold-change = ± 1, and of p-value = 0.05. Box plots demonstrating the percentage of significant clusters identified. Data are shown as mean ± SEM. **(F)** Heatmap showing markers expression of seven clusters. Column scaling was performed to eliminate scale differences between different clusters.

### Unsupervised analysis reveals specific myeloid and immune cell types and distributions in JMML spleens

CD45^+^ immune populations were plotted onto a tSNE immune map that separated key immune lineages, including CD11c^+^/CD11b^+^/CD14^+^myeloid cells, CD123^+^ pDC-like cells, CD56^+^ NK cells, CD19^+^CD20^+^ B cells, CD3^+^CD4^+^ T helper cells, CD3^+^CD8^+^ cytotoxic T cells and CD45^+^CD34^+^ stem cells (Figure 1B). Comparison of samples with somatic or germline *PTPN11* mutation and controls revealed major differences in the immune landscape (Figure 1C).

JMML spleens showed enrichment of stem cells (CD45⁺CD34⁺), pDC-like cells (CD123^+^), and myeloid populations (CD11b⁺ or CD11c⁺ or CD14^+^), accompanied by reduced NK cells, CD4⁺ and CD8⁺ T cells, consistent with manual gating (Figure 1C, Supplementary Figure 1C). PARC clustering identified 35 immune phenotypes across all lineages (Figure 1D, Supplementary Figure 3A). As expected, myeloid subsets were strongly expanded in JMML. B cell populations were also altered in JMML patients, with increased CD73⁺ naïve B cells (c2) and reduced CD27⁺ memory B cells (c5) compared to controls (Figures 1C-D, Supplementary Figure 3B). Noonan syndrome patients with germline *PTPN11* mutations and MPD showed significantly decreased CD11c+ B cells (c18) and CD8⁺ T cells, but expansion of B cell compartment (c6) compared to somatic cases.

Differential expression analysis between controls and JMML samples identified seven strongly differing clusters (Figure 1E). JMML samples showed enrichment of CD123^+^ pDC-like cells (c4), CD45^+^ CD34^+^ CD90^+^ CD33^+^ stem cells (c13), CD11b^+^ CD14^+^ monocytes (c7) and CD56^+^CD16^−^ NK cells (c12), with 3-fold, 5.1-fold, 4.1-fold and 2.4-fold higher abundance, respectively. Interestingly, while c13 stem cells were enriched in JMML, we also identified enriched CD34 stem cells lacking CD45 expression, suggesting expansion of the tumor stem cell compartment (Supplementary Figure 3C). Conversely, CD27^+^ memory B cells (c5), CD4⁺ CD127^+^ CD27^+^ CD45RA^−^ CCR7^−^ effector memory T cells (c14), and CD56^dim^ CD16^bright^ CD11b^dim^ CD27^−^ NK cells (c17), were significantly reduced in JMML samples, with 7.5-fold, 4.7-fold, and 4.8-fold lower abundance, respectively (Figure 1E). Overall, this analysis revealed distinct JMML-associated immune clusters, including expansion of classical monocytes and stem cells.

### Checkpoint molecule expression across immune subsets in JMML cells

We speculated that JMML-altered immune cell types might express targetable immunoregulatory pathways. C7 monocytes expressed several immune checkpoints, including CD39, Siglec-7, Siglec-9, NOX2, and VISTA and CD33, a profile resembling myeloid-derived suppressor cells (MDSCs), known to mediate immunosuppression and immune evasion in malignancies (Figures 1F, 2A-B). pDC-like c4 cells moderately expressed CD47 and NOX2, while c13 stem cells expressed arginase, B7-H3 and CD38, all associated with immunoregulation.

We next assessed the immune checkpoint landscape across all immune lineages in JMML. Myeloid cells expressed multiple immune checkpoints, including CD39, Siglec-7, Siglec-9, CD47, CD38, and NOX2, and lower levels of VISTA, IDO and TIM-3 (Figure 2A). TIM-3 and IDO exhibited moderate expression on dendritic cells, predominantly within clusters c3, c19 and c20. PD-L1 and PD-L2 were weakly expressed, restricted to dendritic clusters c19 and c27. PD-1 and CTLA-4 were mainly expressed on T cells, while CD73 was prominent on B cells (Figure 2A-B). Given the high CD39 expression in the myeloid compartment and the JMML-associated cluster c7, we tested whether CD39 expression was shared across myeloid subsets. CD39 was present in most myeloid cells except pDC-like clusters c4 and c26 (Figure 2C). Overall, JMML immune cells expressed multiple immunoregulatory molecules with potential functional roles, such as promoting adenosine signaling through ectonucleotidases CD39 and CD73.

**Figure 2.**
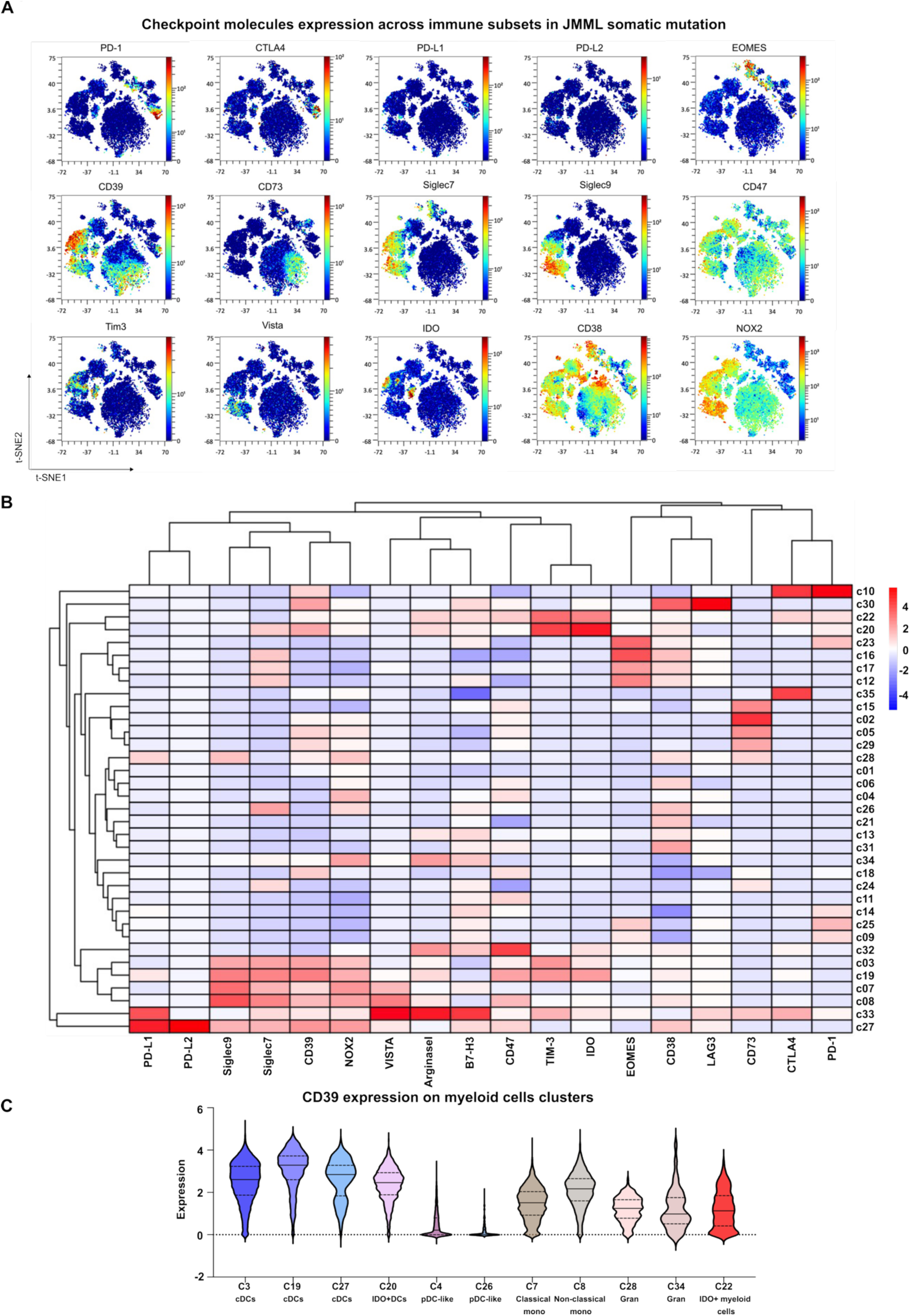
Checkpoint molecule expression across immune subsets in JMML cells. **(A)** Expression of inhibitory molecules on t-SNE. **(B)** Heatmap showing the expression of the inhibitory molecules. Column scaling was performed to eliminate scale differences between different clusters. **(C)** Violin plots of expression of CD39 on myeloid cells clusters in JMML *PTPN11-*mutant samples.

### Murine *Ptpn11^D61Y/+^* positive myeloid cells express various immune escape molecules

To investigate the expression, regulation and function of immune checkpoints downstream of activated SHP2, we used a mouse model expressing oncogenic SHP2^D61Y^ in the hematopoietic compartment, which recapitulates key features of human JMML. In our previous work (22) we defined three distinct stages of progression (Stage 1-3, Figure 3A). Immune escape molecule expression was analyzed in *MxCre;Ptpn11^D61Y/+^*mice 5-7 months after poly(I:C)-induced activation of SHP2^D61Y^ (Figure 3A). At this stage (S2), mice exhibited myeloproliferation and splenomegaly, as seen in JMML, but remained in good overall condition. In addition, we analyzed mice once critically sick (S3). Flow cytometry confirmed significant increase of splenic myeloid populations in S2 and S3 mice (Supplementary Figure 4A), whereas BM myeloid populations were only mildly affected (Supplementary Figure 4B). Further characterization of monocytes (Ly6C^+^Ly6G^−^) and granulocytes (Ly6C^+^Ly6G^+^) revealed significantly increased numbers of cells expressing PD-1, PD-L1, PD-L2, VISTA, CD39 and CD73 in the spleens (Figure 3B-G), which in part was due to the general increase in myeloid cells. However, expression frequency of PD-L1, PD-L2, VISTA, CD39, and CD73 was significantly higher in circulatory monocytes (Figure 3H–M), along with elevated CD39 expression in inflammatory monocytes, indicating a JMML-specific phenotype. Notably, no increased expression of these molecules was observed in BM of *MxCre;Ptpn11^D61Y/+^* mice (Supplementary Figures 4C-H).

**Figure 3:**
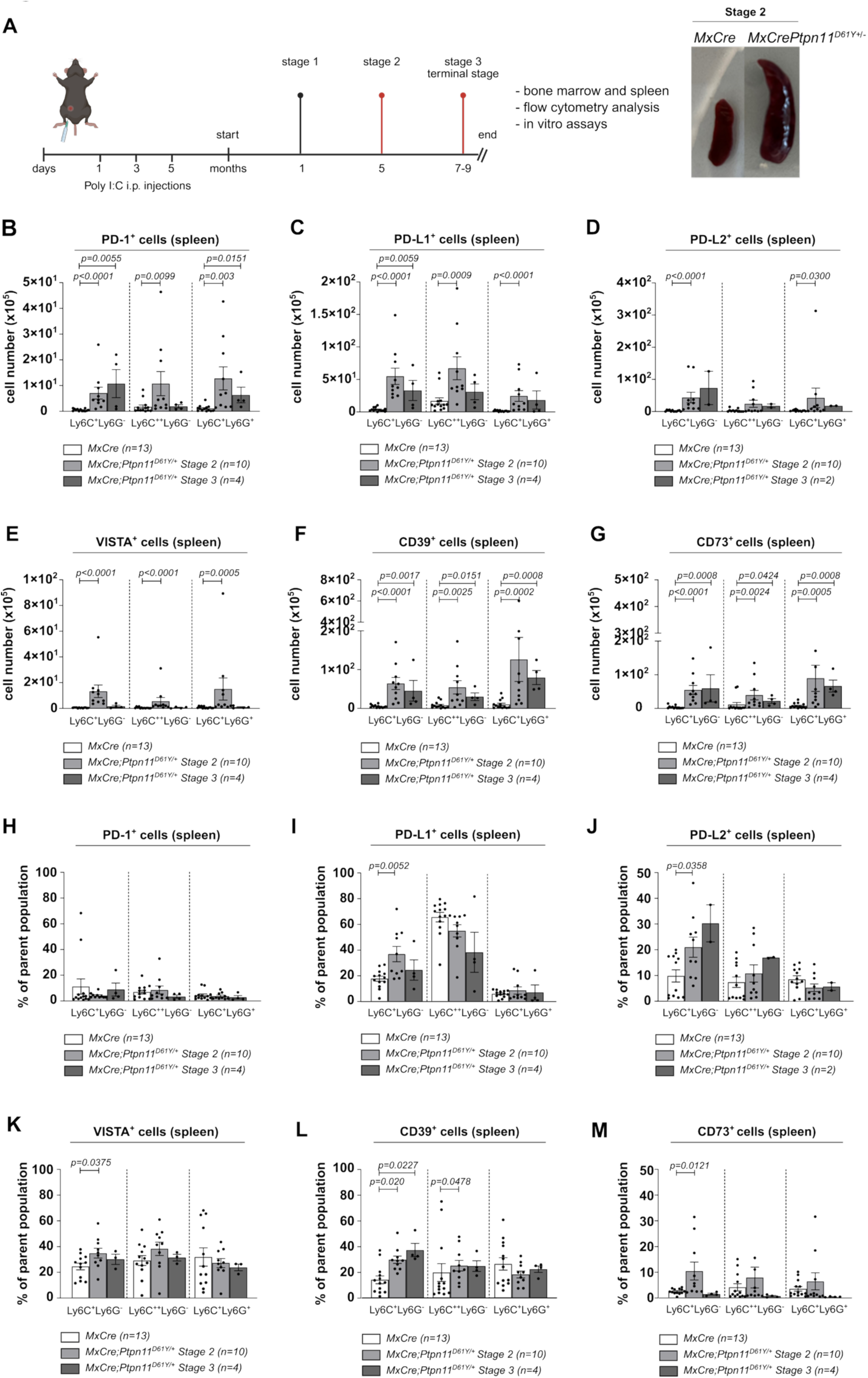
Immune check point molecules are expressed on myeloid cells in *MxCre;Ptpn11^D61Y/+^* mice. **(A)** Schematic illustration of the development of myeloproliferative progression in the mouse model used in this study. *MxCre* (control) and *MxCre;Ptpn11^D61Y/+^* (JMML) mice were intraperitoneally injected with poly(I:C) every other day for three doses and monitored until they reached stage 2 (at 5 months) or stage 3/terminal stage (at 7-9 months) of disease progression. Mice were sacrificed at the indicated time points, and bone marrow and spleen tissues were collected for analysis. **(B-G)** Analysis of PD-1, PD-L1, PD-L2, VISTA, CD39 and CD73 protein levels in myeloid cell subsets from the total spleen cells of the indicated genotypes. **(H-K)** Analysis of PD-1, PD-L1, PD-L2, VISTA, CD39 and CD73 protein on the surface of circulatory (Ly6C^+^Ly6G^−^) and inflammatory monocytes (Ly6C^++^Ly6G^−^) and granulocytes (Ly6C^+^Ly6G^+^) of the indicated genotypes. Data are shown as mean ± SEM from n=4-13 biological replicates per group. Statistical significance was assessed using the Mann-Whitney *U* test.

These findings indicate that SHP2^D61Y^-associated myeloproliferation is accompanied by an immune checkpoint phenotype comparable to that seen in JMML patients.

### JMML monocytes suppress T cell proliferation and activation

To investigate the functional impact of JMML cells on the adaptive immune system, we assessed T cell proliferation and activation under different conditions. First, WT T cells stimulated with anti-CD3/28 beads (Figure 4A-B) were cultured in the presence of serum derived from *MxCre;Ptpn11^D61Y/+^* (S2) or control mice. T cell proliferation remained unaffected irrespective of serum concentration or genotype (Figure 4C).

**Figure 4:**
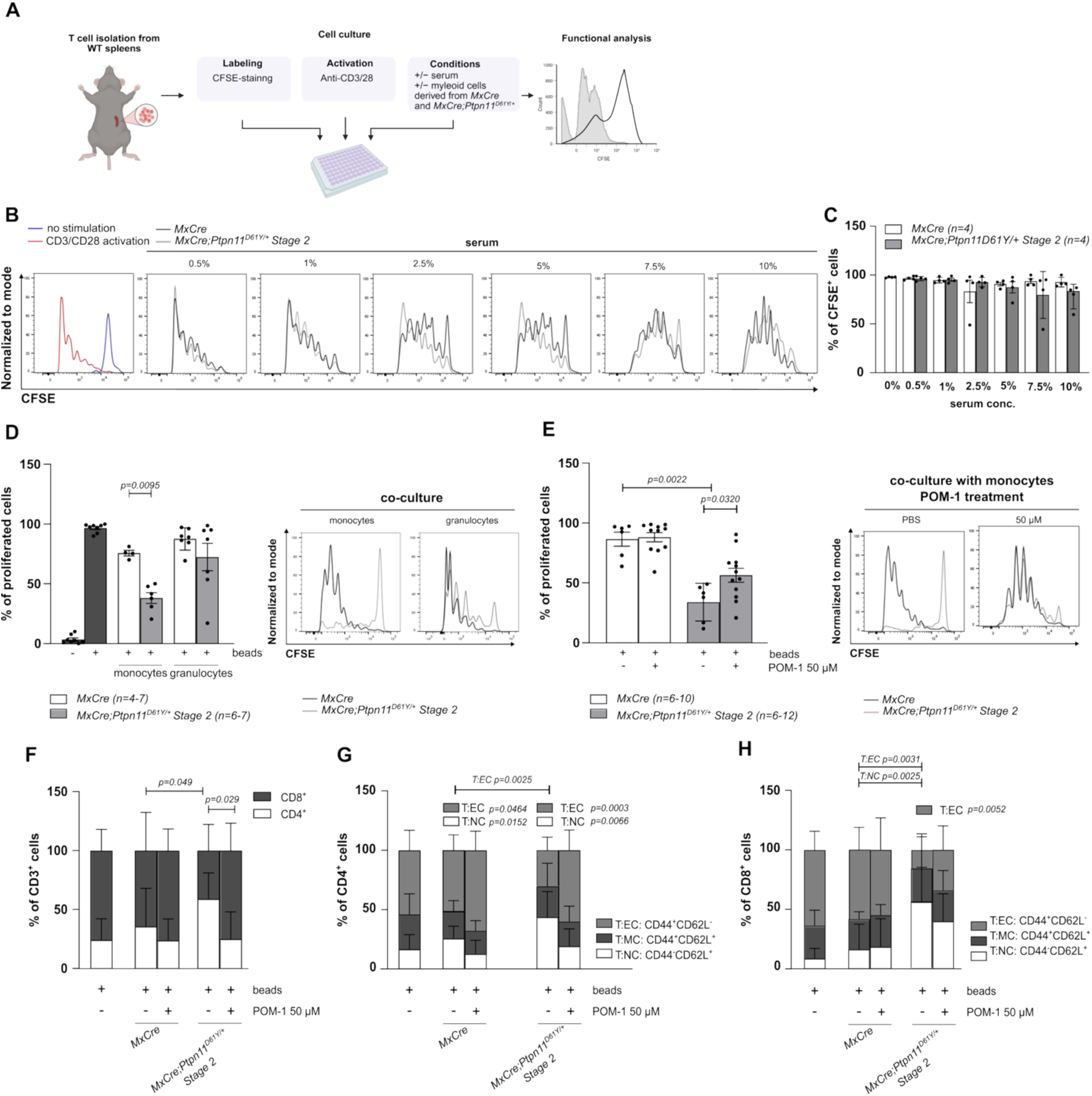
CD39 mediated arrest of T cell proliferation. **(A)** Schematic overview of the experimental setup to assess the influence of *Mx-cre;Ptpn11^D61Y/+^* blood serum, monocytes or granulocytes on T cell function in JMML (created with BioRender). WT T cells were isolated from the spleen, labeled with CFSE, stimulated with CD3/CD28 antibodies, and cultured for 4 days with either blood serum, monocytes or granulocytes isolated from the spleen of *Mx-cre* or *Mx-cre;Ptpn11^D61Y/+^* mice. T cell proliferation was assessed by flow cytometry. **(B)** Representative histogram plots showing CFSE-labeled T cells cultured with serum-supplemented medium. **(C)** Quantification of T cell proliferation in response to serum exposure presented as summary bar graphs. Data are shown as mean ± SEM; n=4 independent experiments. **(D)** Representative histogram plots and summary bar graphs quantifying T cell proliferation in co-culture with FACS-sorted monocytes or granulocytes. Data represent mean ± SEM; n=6–7 independent experiments. **(E)** Representative histograms and bar graphs summarizing T cell proliferation in co-culture with FACS-sorted monocytes treated with POM-1. Data represent mean ± SEM; n=6–12 independent experiments. **(F)** CD4⁺ and CD8⁺ T cell proportions (effector T cells (T:EC: CD44^+^CD62L^−^); memmory T cells (T:MC: CD44^+^CD62L^+^); naïve T cells (T:NC: CD44^−^CD62L^+^ among CFSE-labeled T cells co-cultured with monocytes and treated with POM-1 for 4 days. Data represent mean ± SEM; n=10–12 independent experiments. **(G-H)** Bar graphs showing the distribution of CD4⁺ and CD8⁺ T cells following 4-day co-culture with monocytes and POM-1 treatment. Statistical analysis was performed using the Mann-Whitney test: *P < 0.05, **P < 0.01, ***P < 0.001.

Next, stimulated WT T cells were co-cultured with splenic monocytes or granulocytes from *MxCre* or *MxCre;Ptpn11^D61Y/+^*mice (S2). SHP2^D61Y^ monocytes, but not controls, strongly inhibited T cell proliferation, while granulocytes showed variable and weaker suppressive effects (Figure 4D). To determine whether this suppression required direct cell–cell contact, we performed a transwell assay using spleen-derived monocytes or granulocytes. No inhibitory effect on T cell proliferation was observed (Supplementary Figure 5A), indicating that suppression requires close cell–cell contacts.

We next introduced immune checkpoint inhibitors into the co-cultures. Neither PD-1 blockade nor CD73 inhibition restored proliferation (Supplementary Figures 5B–C). Anti-VISTA showed partial, non-significant rescue, and anti-VISTA/PD-1 combination was ineffective (Supplementary Figure 5E). In contrast, CD39 inhibition with POM-1 almost completely restored proliferation in co-cultures with SHP2-mutated monocytes (Figure 4E). Moreover, POM-1 treatment reversed SHP2^D61Y^ monocyte-induced expansion of CD4^+^ cells and the reduction of effector CD4^+^ and CD8^+^ cells (T:EC; Figure 4F-H). These findings identify CD39 enzymatic activity on SHP2^D61Y^ monocytes as a critical mediator of T cell suppression.

### CD39 and CD73 are downstream of the oncogenic SHP2^D61Y^ signaling

The pronounced effects of POM-1 in co-culture prompted further investigation of CD39 and CD73, ectonucleotidases hydrolyzing extracellular ATP into immunosuppressive adenosine. The adenosine pathway has been implicated immune modulation (25–27). Indeed, the addition of adenosine to WT T cells impeded CD3/CD28-induced proliferation (Supplementary Figure 6A).

We next examined CD39 and CD73 regulation in monocytes and granulocytes from S2 *MxCre;Ptpn11^D61Y/+^* spleens. GM-CSF, a cytokine critically involved in JMML pathogenesis, markedly upregulated CD39 on granulocytes and CD73 on both granulocytes and monocytes. While upregulation of CD39 was similar in *MxCre* and *MxCre;Ptpn11^D61Y/+^* cells, CD73 upregulation was significantly stronger in *MxCre;Ptpn11^D61Y/+^* cells (Figure 5A-B). Inhibition of SHP2/RAS signaling by the MEK-inhibitor trametinib reduced CD39/CD73 on granulocytes and CD73 on monocytes. In granulocytes, CD73 was also regulated by PI3K, another SHP2 target (28), as demonstrated by decreased protein levels upon pictilisib treatment (Figure 5C-F).

**Figure 5:**
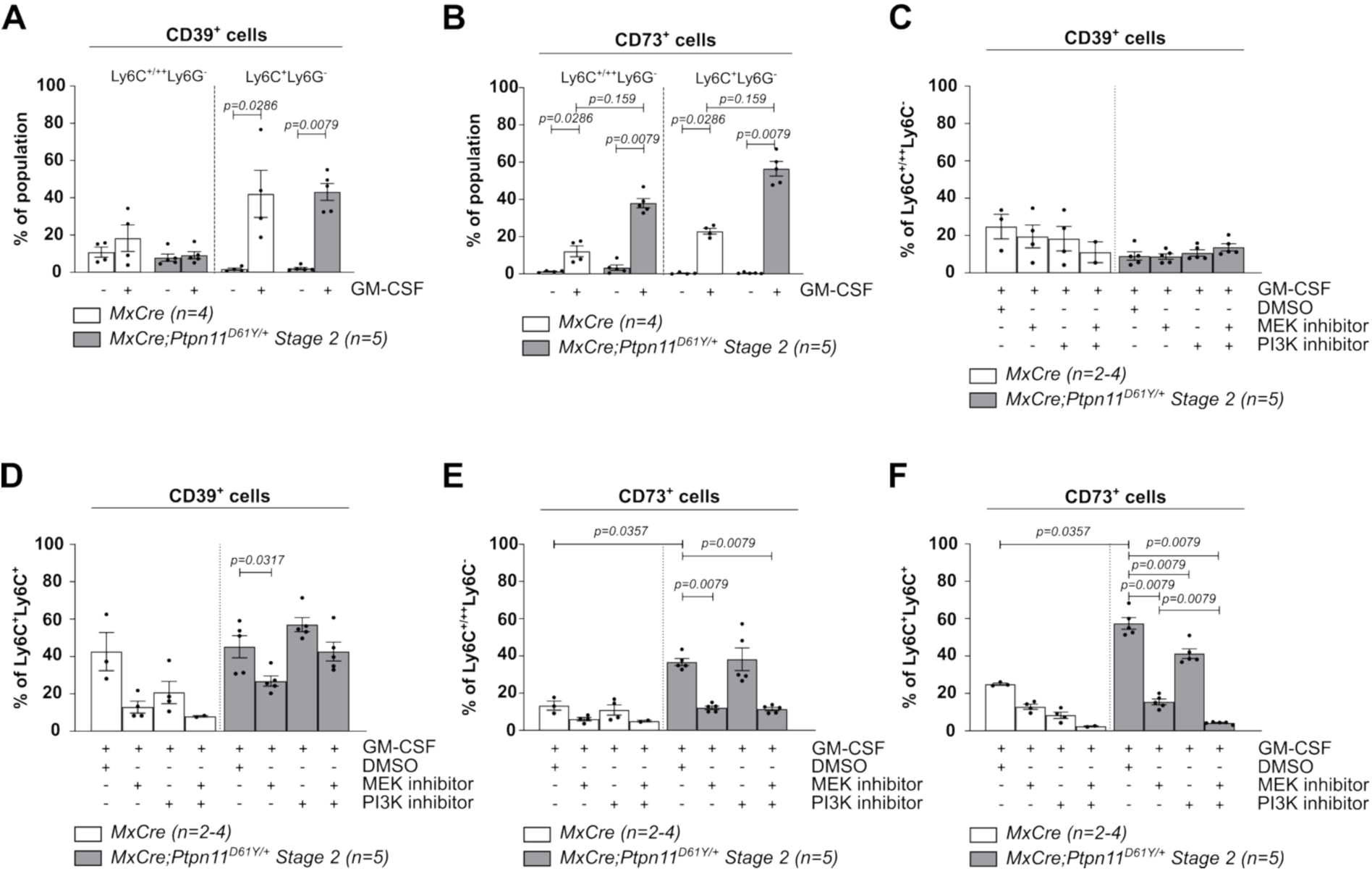
CD39 and CD73 are regulated by the oncogenic SHP2 on myeloid cells in *MxCre;Ptpn11^D61Y/+^*mice. **(A, B)** *In vitro* analysis of CD39 and CD73 expression on monocytes (Ly6C^+/++^Ly6G^−^) and granulocytes (Ly6C^+^Ly6G^+^) derived from the spleen of *MxCre* and *MxCrePtpn11^D61Y/+^.* Cells were cultured in the presence or absence of GM-CSF for 24h. **(C-F)** *In vitro* analysis of CD39 (upper row) and CD73 (lower row) in monocytic (Ly6C^+/++^Ly6G^−^) and granulocytic (Ly6C^+^Ly6G^+^) cell subsets isolated from the spleen of the indicated genotypes. Cells were cultured for 24 hours with GM-CSF and treated with DMSO, MEK inhibitor (trametinib, 1.624 μM), PI3K inhibitor (pictilisib, 50 nM), or a combination of MEK and PI3K inhibitors. Data are presented as mean ± SEM from n=4-5 biological replicates per group. *P* values were determined using the Mann-Whitney *U* test.

To assess whether MEK inhibition also affected other immune checkpoints, we assessed PD-1, PD-L1, PD-L2, and VISTA expression in trametinib-treated myeloid cells from *MxCre;Ptpn11^D61Y/+^* mice and *MxCre* controls. None were elevated in untreated *MxCre;Ptpn11^D61Y/+^* cells relative to controls, nor altered by MEK inhibition (Supplementary Figure 6B-I).

Altogether, these results highlight the CD39/CD73 axis as a distinct and functionally important mediator of JMML-associated immune suppression, regulated by oncogenic signaling and the inflammatory environment.

### *In vivo* CD39 inhibition reduces spleen size and affects immune cell phenotype

To investigate the role of CD39 in myeloid cells, splenic monocytes isolated from *MxCre* and *MxCre;Ptpn11^D61Y/+^*(S2) mice were treated with POM-1 *in vitro*. On one hand, POM-1 effectively inhibited ATP degradation in both genotypes (Figure 6A). Extracellular ATP levels trended to be lower in SHP2^D61Y^ cells treated with POM-1, suggesting slightly increased hydrolysis. On the other hand, POM-1 induced apoptosis in myeloid cells as shown by Annexin V/7AAD staining (Figure 6B).

**Figure 6:**
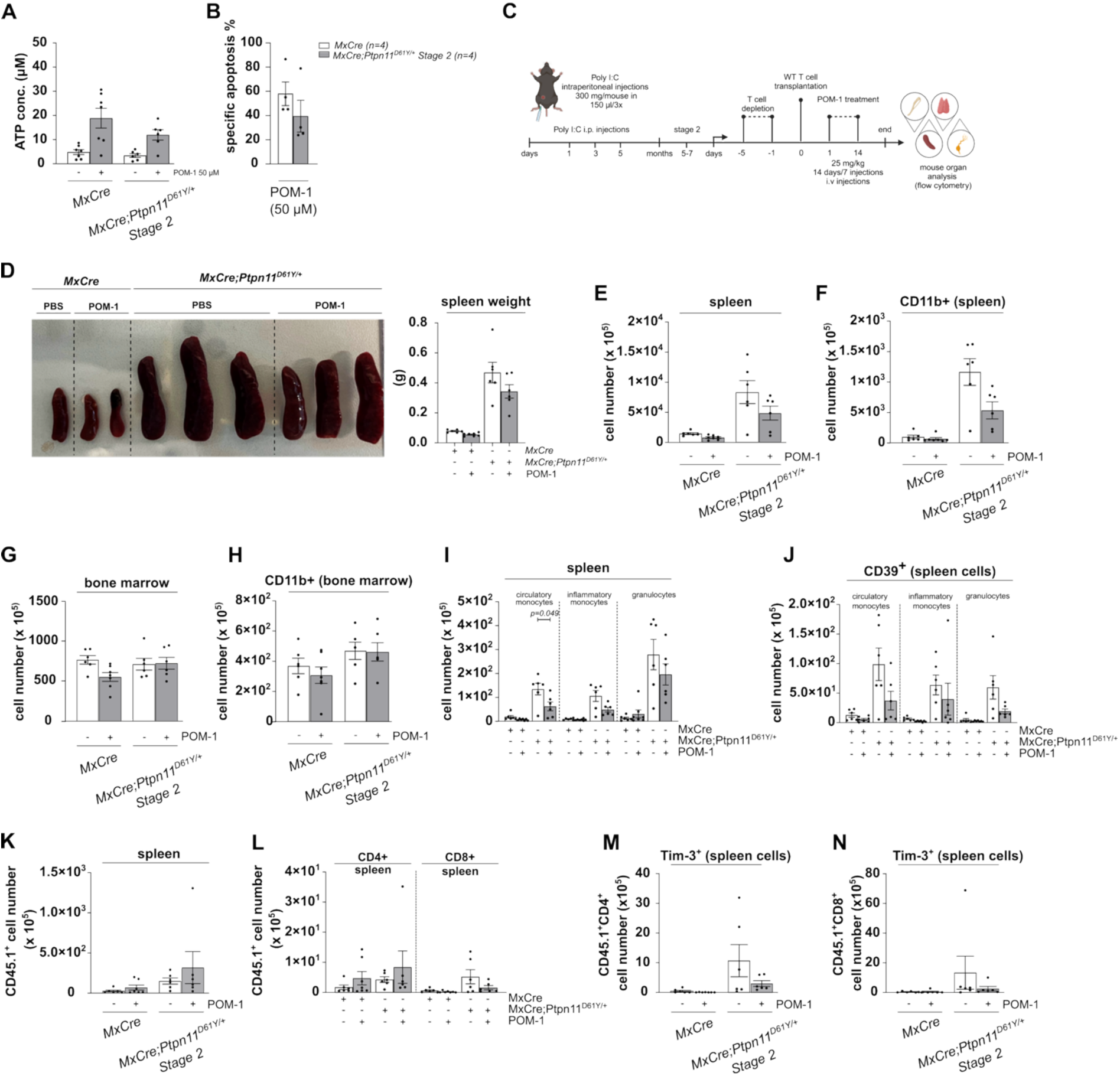
Therapeutic impact of POM-1 on T cell reconstitution and myeloproliferation in *Mx-cre;Ptpn11^D61Y/+^*mice. **(A)** Levels of ATP concentration in the supernatant of *MxCre* and *MxCre;Ptpn11^D61Y^*^/+^ measured by ATP hydrolysis assay. **(B)** The percentage of specific apoptosis induced by POM-1 was determined by flow cytometry using staining for Annexin V and 7AAD. **(C)** Experimental design and therapeutic evaluation of POM-1 in *MxCre;Ptpn11^D61Y/+^* mice. *MxCre* and *MxCre;Ptpn11^D61Y/+^* mice were subjected to *in vivo* depletion of T cell subsets using anti-CD4 and anti-CD8 antibodies. Five days later, mice received adoptive transfer of wild-type (WT) T cells, followed by initiation of POM-1 treatment the next day. **(D)** Clinical evaluation of *MxCre* and *MxCre;Ptpn11^D61Y/+^* mice treated with PBS (vehicle) or POM-1. **(E–F)** Total cell counts in bone marrow and spleen of *MxCre* and *MxCre;Ptpn11^D61Y/+^* mice post-treatment with POM-1. **(G–H)** Quantification of CD11b⁺ myeloid cells in the bone marrow and spleen following POM-1 treatment. **(I)** Distribution of distinct myeloid populations in the spleen of *MxCre* and *MxCre;Ptpn11^D61Y/+^* mice following POM-1 therapy. **(J)** CD39 expression levels across various splenic myeloid subsets. **(K)** Total cell counts of CD45.1⁺ T cells in the spleen following POM-1 treatment. **(L)** Total cell counts of CD45.1⁺CD4⁺ and CD45.1⁺CD8⁺ T cell subsets in the spleen of POM-1 treated and control mice. **(M–N)** Frequency of Tim-3⁺ cells among CD45.1⁺CD4⁺ and CD45.1⁺CD8⁺ T cells in the spleen following POM-1 treatment. Data are presented as mean ± SEM; n=6–7 independent experiments. Statistical significance was assessed using the Mann-Whitney test: *\*p < 0.05, **P < 0.01, ***P < 0.001*.

We next investigated *in vivo* effects of POM-1. Given that *Ptpn11*-mutant T cells are functionally impaired (29), our model was optimized by autologous T-cell depletion and adoptive transfer of 5 × 10⁶ congenic (CD45.1) WT T cells into *MxCrePtpn11^D61Y/+^* mice, followed by POM-1 or vehicle (PBS) injection every other day for 14 days (Figure 6C, Supplementary Figure 7A).

In POM-1 treated mice we observed a reduction in spleen size in *MxCre;Ptpn11^D61Y/+^* mice, in line with decreased splenic cellularity (Figure 6D-E) and the frequency of myeloid cells (Figure 6F; Supplementary Figure 7B). In contrast, BM cells were not affected (Figure 6G-H; Suppl. Figure 7C). Circulating monocytes were significantly reduced in spleens of *MxCre;Ptpn11^D61Y/+^*mice, while inflammatory monocytes and granulocytes were only moderately reduced (Figure 6I). POM-1 preferentially depleted CD39⁺ cells (Figure 6J).

Transferred CD45.1⁺ T cells were detected at low levels in spleens of both genotypes but modestly increased after POM-1 treatment (Figure 6K; Supplementary Figure 7D). In controls, but not *MxCrePtpn11^D61Y/+^* mice, POM-1 slightly elevated splenic CD4⁺ T cells, while CD8⁺ T cells were slightly reduced upon treatment (Figure 6L; Supplementary Figure 7E). In *MxCrePtpn11^D61Y/+^* spleens, POM-1 reduced CD4⁺TIM-3⁺ and CD8⁺TIM-3⁺ T subsets, suggesting partial reversal of immune exhaustion (Figure 6M-N).

To determine systemic effects, plasma ATP and metabolites were measured by LC-MS/MS. ATP, ADP, AMP, and adenosine levels were comparable between genotypes, both at baseline and after POM-1 treatment. This indicates that neither myeloproliferation nor POM-1 treatment affected systemic levels of these nucleotides (Supplementary Figure 6F-I).

Taken together, CD39 inhibition exerts *in vitro* and *in vivo* effects on the myeloid and the adaptive immune compartment of *MxCre;Ptpn11^D61Y/+^*mice.

## DISCUSSION

With the aim of developing novel and effective therapeutic strategies for JMML patients, and in light of the therapeutic relevance of immune escape in myeloid malignancies like AML (12), our study investigated the immune escape profile in JMML with *PTPN11* mutations.

High-dimensional immune profiling of *PTPN11*-mutant JMML patient spleens revealed profound alterations in both innate and adaptive immune compartments. JMML samples exhibited expansion of stem and progenitor cells, MDSC-like populations and pDC-like cells, alongside a reduction in NK and T cells. These findings indicate that JMML is associated with a coordinated remodeling of the splenic microenvironment. Notably, SHP2-activating mutations in mice result in similar accumulation of HSCs and myeloid cell types, in part caused by increased cell survival through upregulation of anti-apoptotic proteins such as BCL-XL and MCL-1 (22, 30). Importantly and in line with the indolent clinical presentation, germline *PTPN11*-mutant Noonan syndrome patients with MPD displayed distinct immune profiles in their spleens.

The immune checkpoint landscape revealed that JMML-associated myeloid cells express multiple immunoregulatory molecules, including CD39, Siglec-7, Siglec-9, NOX2, VISTA, and CD33, resembling MDSC-like phenotypes observed in different tumor types (31–33). This immunosuppressive profile supports the concept that myeloid cells belonging to the JMML clone contribute to immune evasion. Unlike many solid tumors where immune evasion commonly relies on T-cell checkpoints such as PD-1 and CTLA-4 engaged by their ligands (34), JMML shows only weak expression of PD-L1 and PD-L2.

Our murine *MxCre;Ptpn11^D61Y/+^* model recapitulated findings in human JMML samples, demonstrating elevated expression of various immune checkpoints including CD39, CD73, VISTA, PD-1, PD-L1 and PD-L2, in clonal monocytes and granulocytes. Functional assays confirmed that SHP2^D61Y^-expressing monocytes, and to a more variable degree also granulocytes, potently suppressed T cell proliferation and effector differentiation via direct cell– cell interactions, with CD39 identified as the principal mediator. CD39 has been increasingly recognized as an important factor in shaping the immunosuppressive tumor microenvironment (35, 36). It can be involved in mediating a functional exhaustion of CD8+ T cells that precedes PD-1 expression (37). Both, CD39 and CD73, ectonucleotidases responsible for converting ATP to adenosine, have been shown to allow tumor cells to escape cytotoxic T cell–mediated immunity in human lung cancer and in a mouse model of ovarian cancer (33, 38).

CD39’s enzymatic product, adenosine, is an established immunosuppressive metabolite in solid tumors, and similar findings in AML further support its role in promoting an immunosuppressive microenvironment (39–41). Conversely, blocking CD39 reduces ATP hydrolysis, leading to accumulation of extracellular ATP (eATP), which can exert direct cytotoxic effects on tumor cells. Indeed, eATP has been reported to reduce leukemia growth *in vivo* and to enhance the antileukemic activity of cytarabine in AML (40). Our findings extend these observations to JMML. Moreover, the adenosine axis appears more pivotal than classical PD-1/PD-L1 inhibitory pathway, consistent with the lower PD-L1/PD-L2 expression observed in human samples. Indeed, treatment of murine SHP2^D61Y^-expressing monocytes with a PD-1-blocking antibody did not rescue T cell proliferation in the co-culture experiments. We found that GM-CSF, a cytokine implicated in JMML pathogenesis (42), enhanced CD39 and CD73 expression on SHP2^D61Y^-expressing granulocytes, as well as CD73 on SHP2^D61Y^-expressing monocytes. CD39 was regulated through the MAPK signaling (41), whereas CD73 was regulated by both MAPK and PI3K signaling downstream of SHP2. These findings highlight that the oncogenic SHP2 signaling directly affects the immune escape phenotype of clonal myeloid cells.

A direct connection between oncogenic SHP2 signaling and immune evasion has previously been demonstrated in other cancer types. *In vitro* inhibition of SHP2 sensitizes cancer cells to IFNγ, thereby promoting T cell proliferation, activation, and cytotoxicity in ovarian spheroids, while *in vivo* it amplifies tumor IFNγ signaling, strengthens cytotoxic T cell function, reduces immunosuppressive myeloid cells, and ultimately enhances tumor control across breast and colon cancer models (21). In melanoma, myeloid-restricted SHP2 deletion suppresses tumor growth by increasing macrophage-derived CXCL9 and reinforcing a CXCL9–IFNγ–T cell feedback loop, thereby supporting antitumor T cell responses (43). These findings illustrate that SHP2 drives immune escape through mechanisms that are highly dependent on the tumor entity and cell type. In our murine JMML model, only CD39 and CD73, but not PD1, PD-L1, PD-L2, or VISTA, were directly regulated by oncogenic SHP2.

*In vivo*, POM-1 partially reduced splenic cellularity, depleted CD39⁺ myeloid cells, and modestly reduced Tim3^+^ T subsets indicating local remodeling of the splenic immune environment. Supporting the broader relevance of CD39 targeting, preclinical melanoma studies have shown that POM-1 reduces metastatic burden and enhances antitumor immunity, and CD39-deficient mice similarly display resistance to metastasis (44). The emerging importance of CD39 in regulating the tumor immune microenvironment has spurred the development of targeted therapies, with several CD39-blocking antibodies, including TTX-030, IPH5201, and SRF617, currently in clinical trials for solid tumors and lymphomas, either as monotherapies or combined with chemo-or immunotherapy (45–47). These findings collectively highlight CD39 as a candidate therapeutic target in JMML – albeit not as monotherapy. In addition to the immune-modulatory effects, POM-1 induced apoptosis in SHP2-mutant myeloid cells, demonstrating a cell-intrinsic function of CD39 for JMML cells.

Together, these findings identify CD39 as a central mediator of immune evasion in JMML and highlight it as a promising therapeutic target.

## AUTHORSHIP CONTRIBUTION

J.R., K.M., B.B. and M.E. designed, interpreted and analyzed experiments and wrote the manuscript; J.R, K.M., J.W., A.E.S., S.B. and B.B. performed and analyzed the CyTOF; L.H. performed metabolite mass spectrometry; J.R., K.M., A.L.S., J.W., N.K., H.X. and L.B. performed experiments; R.Z and D.B.L. helped in concept development and data interpretation; C.N., C.F., R.M. and M.E. took care of the patients and provided patient material.

## ACKNOWLEDGMENTS

The authors thank U. Kern for the valuable discussions and N. Kaltenbach and S. Ehret for excellent technical support. They also acknowledge N. Krause and her team at the Center for Experimental Models and Transgenic Services for their dedicated care of the animals. Special thanks go to M. Follo and her team at the Lighthouse Fluorescence Technologies Core Facility, Freiburg, for expert assistance with cell sorting, and the members of the Freiburg mass cytometry Facility. We thank A. Büttgenbach and A. Mesesan for help with CyTOF experiments. We are grateful to the Shared Translational Metabolomics Core Facility, MetaboCF, Faculty of Medicine, Medical Center – University of Freiburg, Germany for providing support and instrumentation, Deutsche Forschungsgemeinschaft (DFG, German Research Foundation) – Research Infrastructure ID: RI_00507, and Projektnummer 2023/A7-Han. The authors acknowledge the services of the Hilda Biobank for Children at the Children’s Hospital Freiburg, Germany. Studies EWOG-MDS 96 and 2006 were supported by the parent association “Förderverein für krebskranke Kinder e.V. Freiburg, Germany”. J.R. and K.M. received support from the Oncoescape Graduate School, with additional support for J.R. provided by the Spemann Graduate School of Biology and Medicine at the University of Freiburg. K.M, J.W and H.X were supported by Chinese Government Council (CSC) scholarship. This project was performed in the framework of the Collaborative Research Center (CRC) 1479 Oncoescape (grant no. 441891347 – P16 to ME and BB, P01 to RZ). M.E. received funding from the German Federal Ministry of Education and Research (BMBF), Berlin, through the project MyPred – Network for Young Individuals with Syndromes Predisposing to Myeloid Malignancies (grant no. 01GM2207A), as well as from the DFG under the framework of TRR 353/1 – Regulation of Cell Death Decisions (grant no. 471011418 – A06) and José Carreras Leukämie -Stiftung (grant no. DJCLS 10R/2019).

B.B. was further supported by the DFG under the frameworks of Heisenberg Program (grant no. 520992132), Center for Integrative Biological Signaling Studies (CIBSS, grant no. 390939984), TRR179 (grant no. 72983813), FOR5560 (grant no. 505372148) and the German Cancer Consortium (DKTK).

---

**Supplementary Figure 1.**
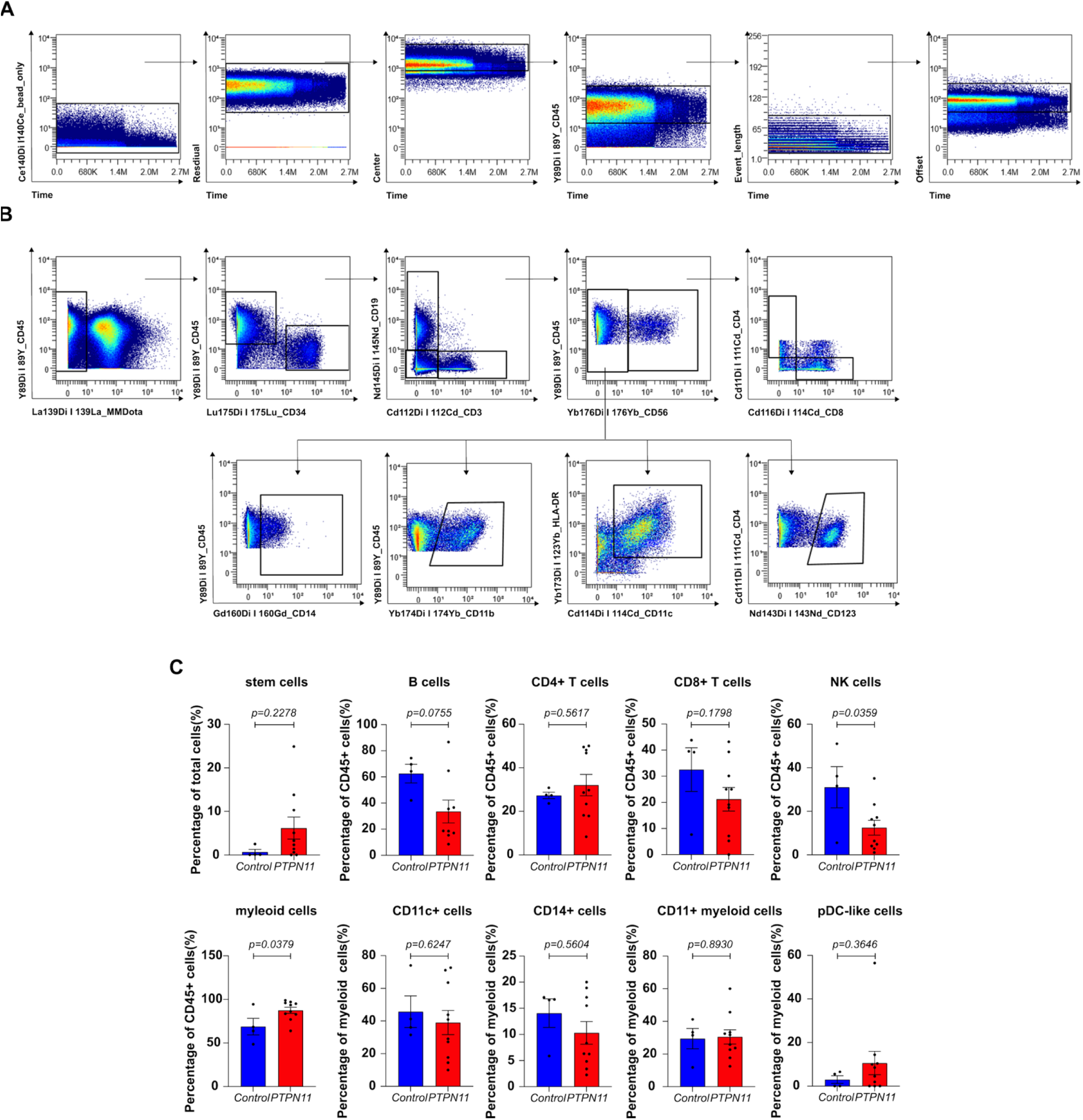
Preprocessing workflow, manual gating, and statistical comparison of splenic immune cells in JMML *PTPN11*-mutant and control samples. **(A)** Strategies for data preprocessing before high-dimensional data reduction analysis. **(B)** Manual gating on splenocytic live cells in JMML *PTPN11*-mutant and control samples. **(C)** Statistical analysis of manually gating on splenocytic live cells in JMML *PTPN11*-mutant (n=10) and control samples (n=4) by paired two-tailed Student’s t-test for normally distributed data or Mann-Whitney test for non-normally distributed data. Data are shown as mean ± SEM.

**Supplementary Figure 2.**
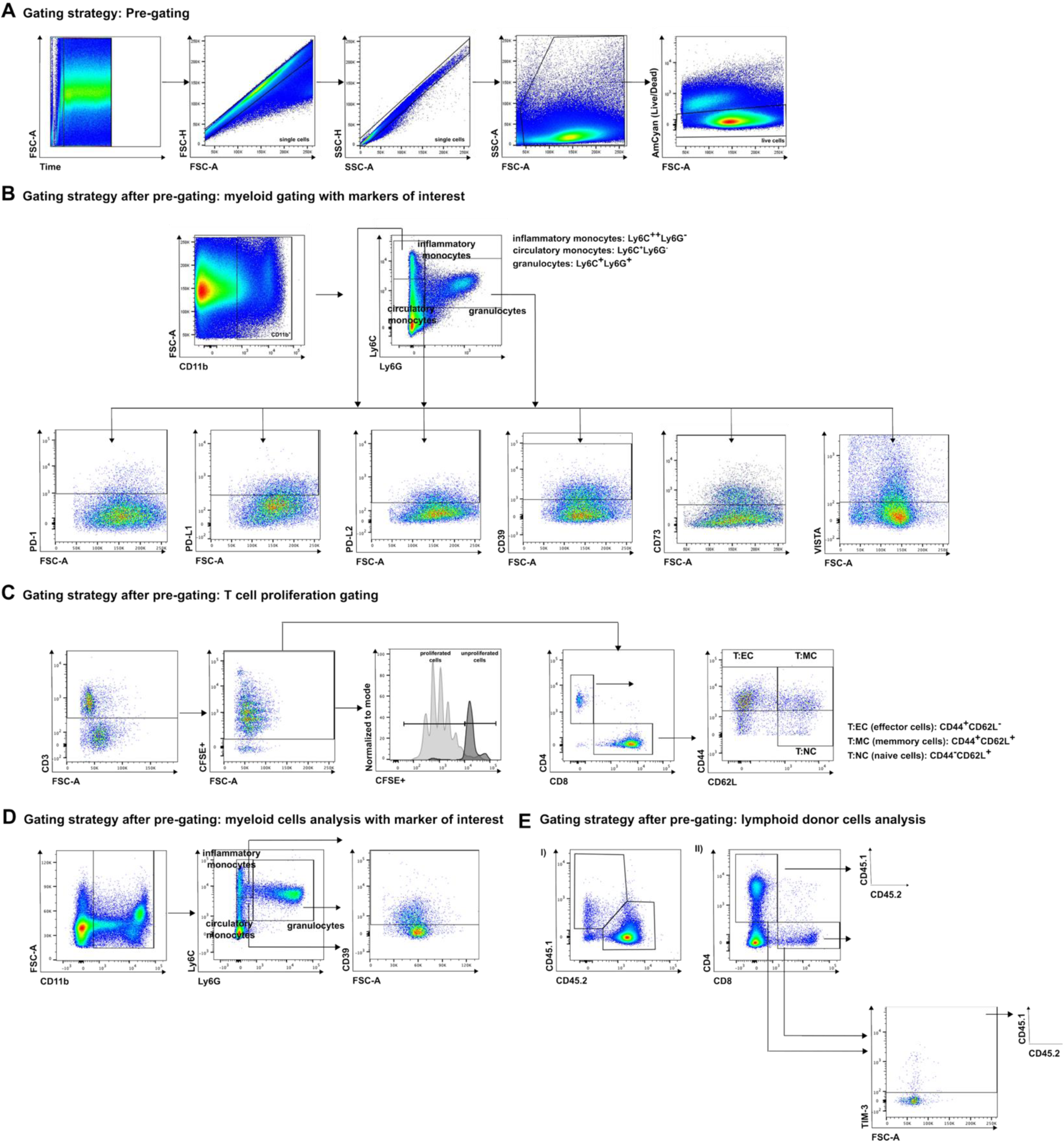
Flow cytometry gating strategy. **(A)** Overview of data preprocessing steps prior to analysis. **(B)** Gating strategy for identifying myeloid cell subsets and immune checkpoint markers in BM and spleen. **(C)** Gating of proliferating T cells and delineation of distinct T cell subsets. **(D)** Gating strategy for myeloid cell populations and CD39 expression in BM and spleen. **(E)** (I) Gating of transplanted wild-type (WT) CD45.1⁺ cells in BM and spleen. (II) Gating of CD4⁺, CD8⁺, and TIM3⁺ T cells within the transplanted WT CD45.1⁺ population.

**Supplementary Figure 3.**
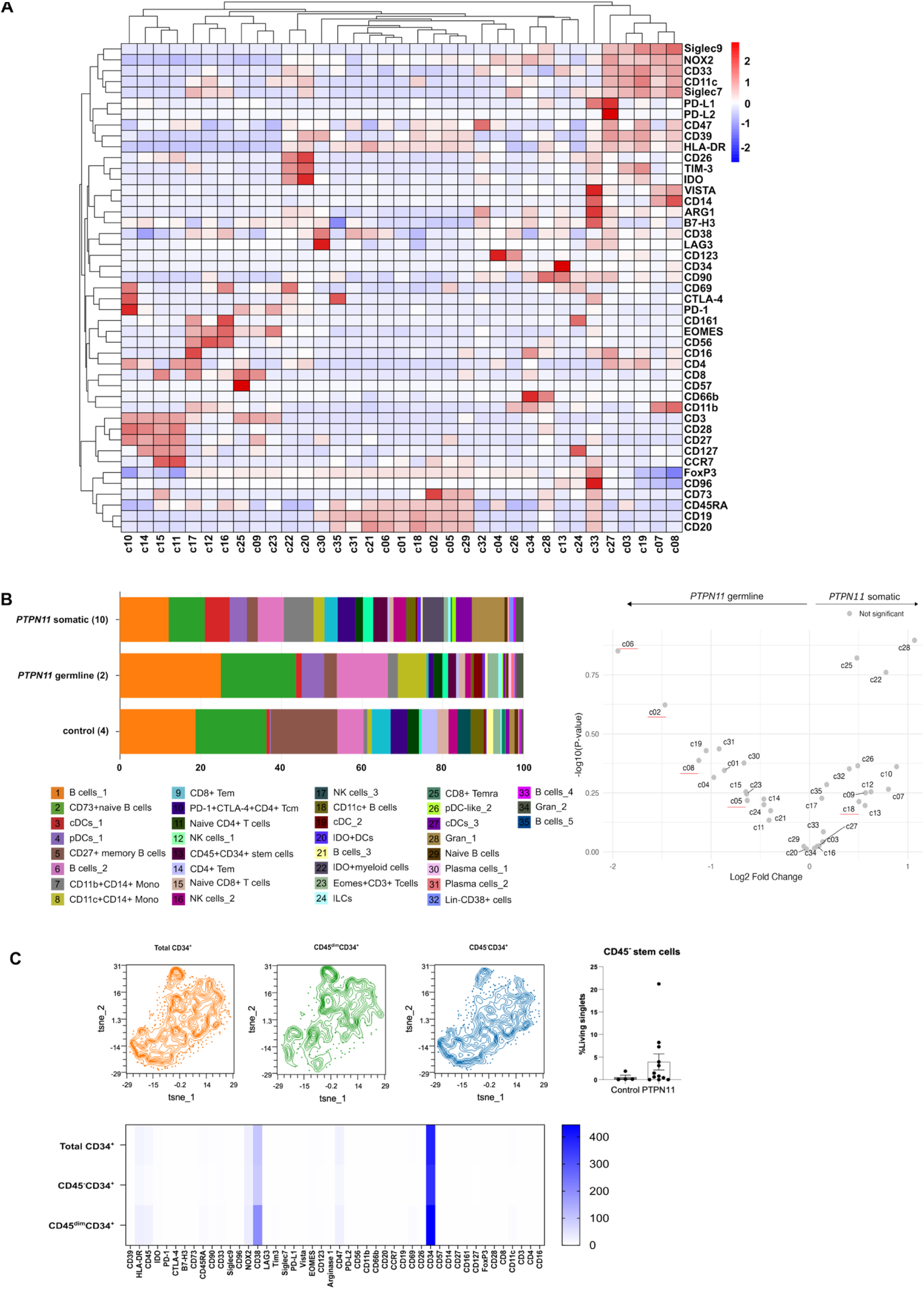
Comprehensive profiling of cells clusters, differential analysis across groups and Characterization of hematopoietic stem cell subpopulations. **(A)** Heatmap showing the expression of 45 markers. Row scaling was performed to eliminate scale differences between different clusters. **(B)** Barplots of Cluster Frequencies in Somatic, Germline, and Control Samples. Volcano plots illustrating the differential expression analysis between germline mutation group and somatic mutation group. Each point represents a cluster, with the y-axis showing the fold change (log2 scale) and the x-axis representing the statistical significance (-log10 p-value). Significant clusters are highlighted in orange (upregulated) or blue (downregulated) above the cut-off of fold-change = ± 1, and of p-value = 0.05. **(C)** Contour plots showing the cells distribition among the following three group: total CD34^+^, CD45^−^CD34^+^ and CD45dim CD34^+^. Heatmap showing that the expression levels of immune inhibitory/immune checkpoints molecules and differentiation/stemness markers among these three group. Box plot demonstrating the counts of CD45^−^CD34^+^ cells between Control and *PTPN11* group. Data are shown as mean ± SEM.

**Supplementary Figure 4:**
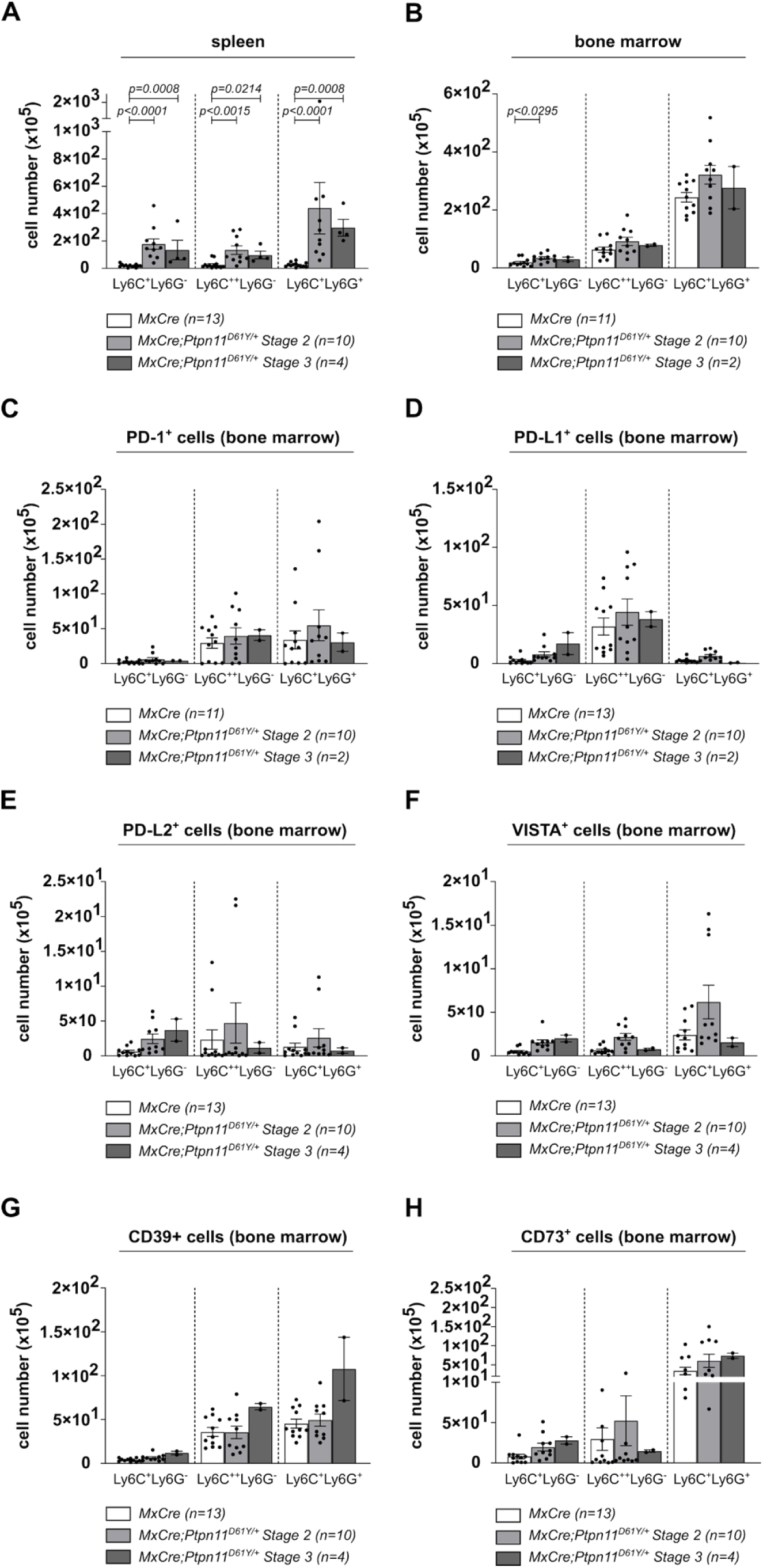
Immune check point molecules expression on myeloid cells derived from bone marrow of *MxCre;Ptpn11^D61Y/+^* mice. **(A-B)** Total number of circulatory and inflammatory monocytes and granulocytes in the spleen and bone marrow of the indicated genotypes and disease stages. Data are shown as mean ± SEM from n=2-13 biological replicates per group. Statistical significance was assessed using the Mann-Whitney *U* test. **(C-H)** Analysis of PD-1, PD-L1, PD-L2, VISTA, CD39 and CD73 protein expression in circulatory (Ly6C^+^Ly6G^−^) and inflammatory monocytes (Ly6C^++^Ly6G^−^) and granulocytes (Ly6C^+^Ly6G^+^) from the bone marrow of the indicated genotypes. Data are shown as mean ± SEM from n=4-13 biological replicates per group. Statistical significance was assessed using the Mann-Whitney *U* test.

**Supplementary Figure 5:**
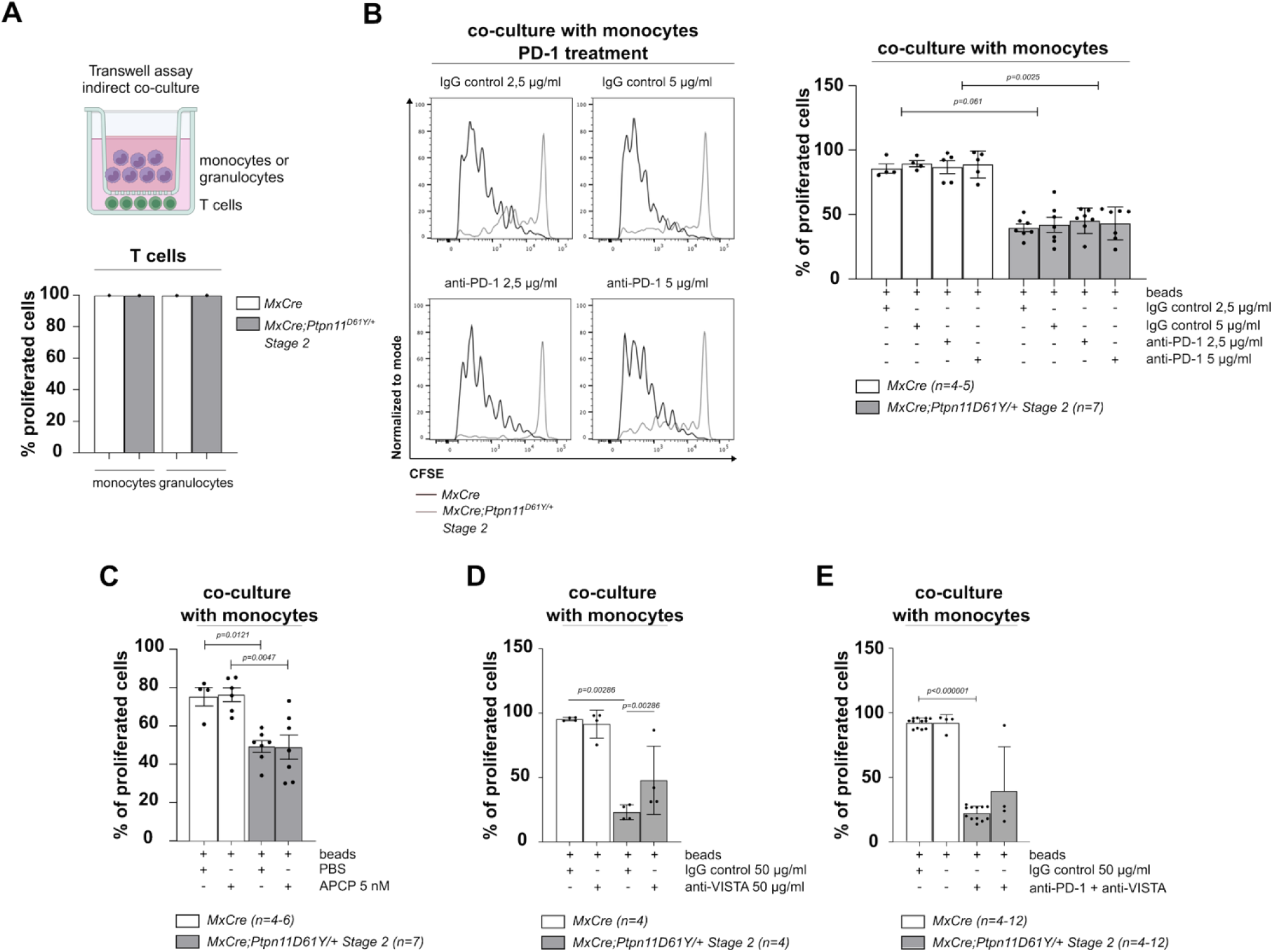
Assessment of T cell proliferation in transwell and co-culture assays with myeloid cells and immune checkpoint inhibitors. **(A)** Schematic representation of the experimental design (created with BioRender). Briefly, CFSE-labeled WT T cells were isolated from spleen, stimulated with CD3/CD28 antibodies, and cultured for 4 days in a transwell assay in the presence of either monocytes or granulocytes assessed by flow cytometry and shown as bar graphs. **(B)** Representative histogram plots and bar graphs showing CFSE-labeled T cells co-cultured with FACS-sorted monocytes for 4 days and treated with anti–PD-1 antibody or isotype IgG control. Data are shown as mean ± SEM; n=4–12 from independent experiments. **(C)** Bar graphs quantifying T cell proliferation after co-culture with FACS-sorted monocytes for 4 days under PBS or APCP treatment. Data are shown as mean ± SEM; n=4–6 from independent experiments. **(D)** Bar graphs quantifying T cell proliferation after co-culture with FACS-sorted monocytes for 4 days under anti–VISTA or IgG control treatment. Data are shown as mean ± SEM; n=4–5 from independent experiments. **(E)** Bar graphs quantifying T cell proliferation after co-culture with FACS-sorted monocytes for 4 days under anti–anti-PD-1 and VISTA or IgG control treatment. Data are shown as mean ± SEM; n = 4–12 from independent experiments. Statistical analysis was performed using the Mann-Whitney test: *P < 0.05, **P < 0.01, ***P < 0.001.

**Supplementary Figure 6:**
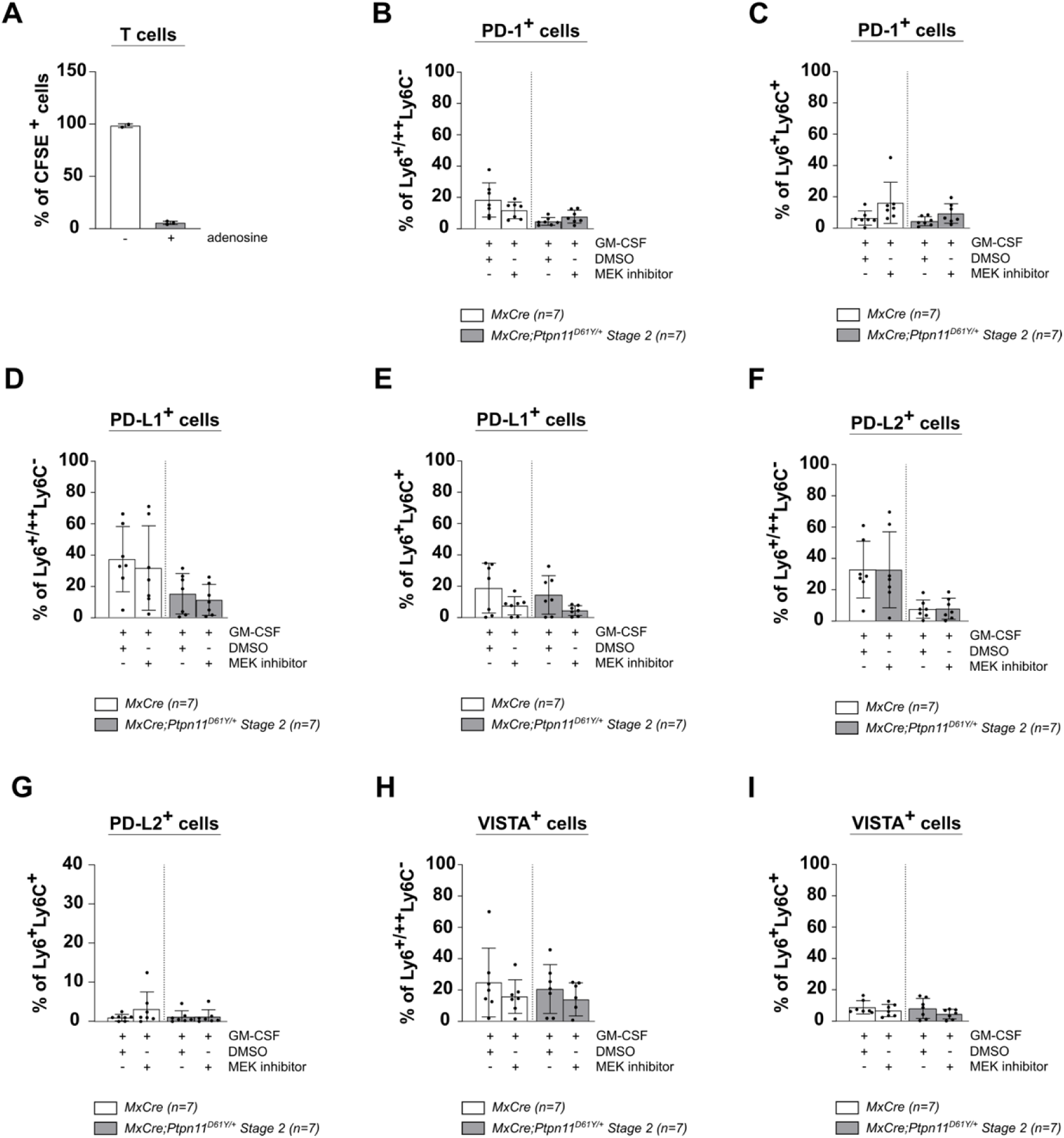
Expression and regulation of immune checkpoint molecules on myeloid cells in *MxCre;Ptpn11^D61Y/+^* mice. **(A)** WT T cell proliferation in the presence of adenosine (100 µM) was assessed using flow cytometry analysis after 96 h. **(B-I)** *In vitro* analysis of PD-1, PD-L1, PD-L2 and VISTA expression in monocytic and granulocytic cell subsets isolated from spleen of the indicated genotypes. Cells were cultured for 24 hours with GM-CSF and treated with DMSO or MEK inhibitor (trametinib, 1.624 μM). Data are presented as mean ± SEM from n=7 biological replicates per group. *P* values were determined using the Mann-Whitney *U* test.

**Supplementary Figure 7:**
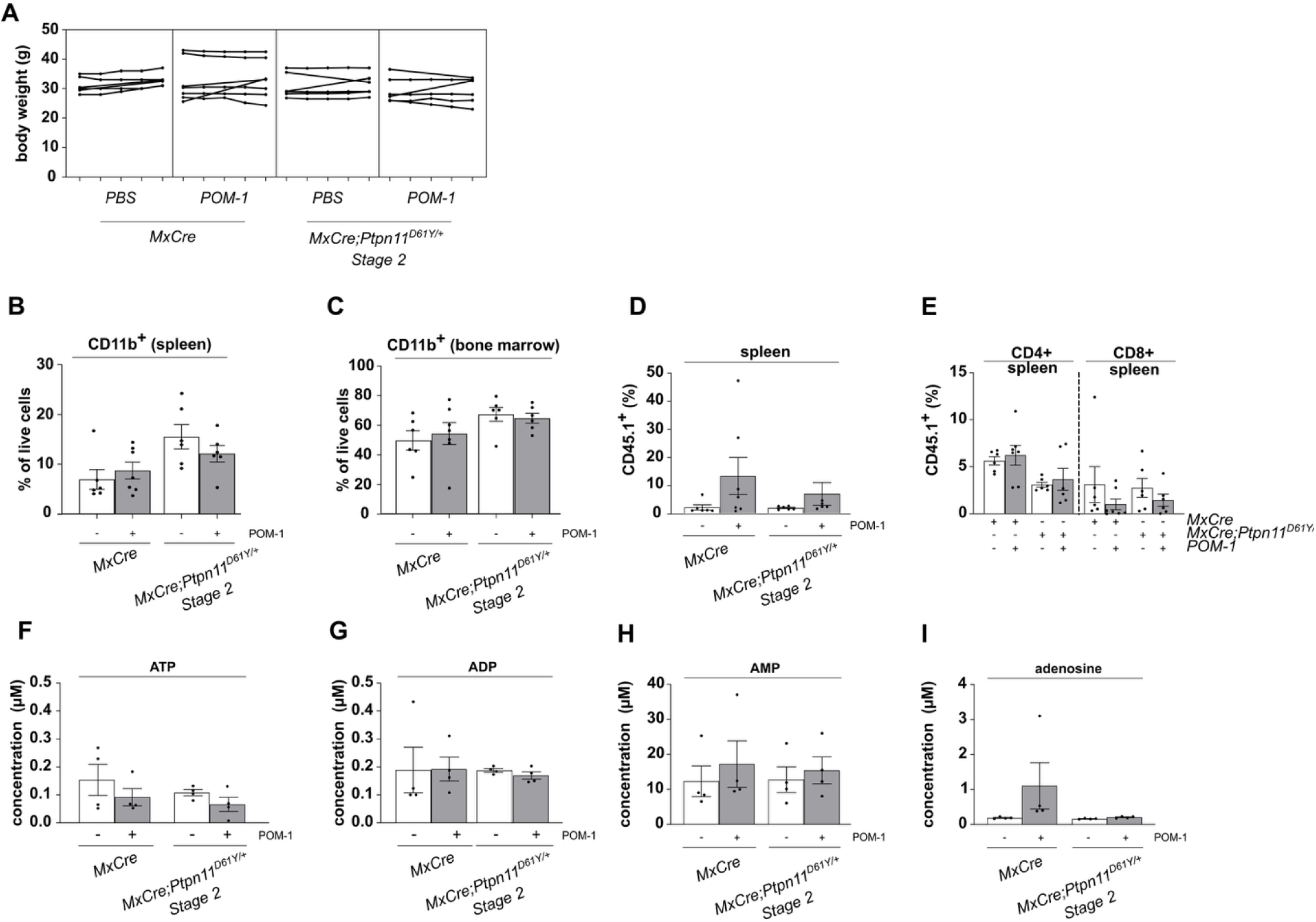
POM-1–induced changes in immune cell dynamics and purine metabolism. **(A)** Body weight changes in *MxCre* and *MxCre;Ptpn11^D61Y/+^* mice throughout the course of POM-1 treatment. **(B–C)** Frequency of CD11b⁺ myeloid cells in the bone marrow and spleen of *MxCre* and *MxCre;Ptpn11^D61Y/+^* mice following POM-1 administration. **(D)** Proportion of CD45.1⁺ T cells in the spleen post POM-1 treatment. **(E)** Frequency of CD45.1⁺CD4⁺ and CD45.1⁺CD8⁺ T cell subsets in the spleens of POM-1-treated and control mice. **(F–I)** Quantification of ATP, ADP, AMP, and adenosine, in the plasma of *MxCre* and *MxCre;Ptpn11^D61Y/+^* mice after treatment with POM-1. Data are presented as mean ± SEM; n=6– 7 independent experiments. Statistical analysis was performed using the Mann-Whitney test: *\*p < 0.05, **P < 0.01, ***P < 0.001*.

