## Supplementary Methods for "Oncogenic PTPN11/SHP2 drives immune escape in juvenile myelomonocytic leukemia (JMML) through activation of ectonucleotidase/adenosine signaling"

**Mass Cytometry**

**Preparation of single-cell suspensions:** Clumping cells were carefully dissociated using Accumax at 37 °C with intermittent vortexing, filtered through a 70 µm strainer, and combined with the main suspension.

**Staining procedure:** Antibodies were either pre-conjugated from Fluidigm or conjugated in-house using Maxpar X8 or MCP9 Antibody Labeling Kits (Fluidigm). Metal-labeled antibody titration was performed on peripheral blood mononuclear cells (PBMCs) from healthy donors. The washes used DPBS (PAN BIOTECH) containing 0.1% FCS (Fisher Scientific). Fixation was performed with 1.6% paraformaldehyde (Invitrogen) for 5 minutes at RT. Cells were permeabilized using the eBioscience™ Foxp3/Transcription Factor Staining Buffer Set (Invitrogen) at 4°C for 30 minutes and washed twice with FoxP3 buffer (Invitrogen) before intracellular staining.

**Data Acquisition and Analysis** A spillover compensation matrix was generated using CATALYST (24). Expression values were transformed using an inverse hyperbolic sine (arcsinh) transform with cofactor 5. Sequential gating included “Time vs Beads,” “Time vs Residual,” “Time vs Center,” “Time vs Offset,” “Time vs Event Length,” and “Time vs CD45.” Subsampling was applied (max 10,000 events per sample), with all markers included for t-SNE analysis. Parc clustering was applied to all CD45^+^ living cells. For stem cell subsets, t-SNE used all markers except lineage markers. (Supplementary Figure 1A-B).

***In vivo* treatment with the CD39 inhibitor POM-1**

Approximately 25 weeks after poly(I:C) treatment, endogenous *Ptpn11^D61Y/+^* T cells were depleted via i.p. injections of anti-CD4 (250 µg; clone GK1.5; ichorbio) and anti-CD8 (250 µg; clone YTS-169; ichorbio) antibodies in 150 µL PBS on days -8 and -4. On day 0, mice received 5×10⁶ CD45.1⁺ T cells via tail vein. POM-1 (Tocris) was reconstituted in sterile water, diluted in PBS, and administered i.v. PBS served as vehicle control. Gating strategy for flow cytometry is shown in Supplementary Figure 2.

**Quantification of adenosine, AMP, ADP and ATP in murine** **plasma**

Gradient: 0–0.5 min: 2% B; 0.5–1.5 min: 2% B; 1.5–3.5 min: 100% B; 3.5–5.5 min: 2% B. Retention times (min): adenosine 3.1, AMP 1.03, ADP 1.02, ATP 1.05, 15N5-ATP 0.95. Method validation included analysis of healthy mice and human plasma and comparison to Human Metabolome Database values. Mass spectrometry settings are in Supplementary Table 5.

**Myeloid cells isolation and culture**

Cells were sorted using FACSAria Fusion, FACSAria III, or MoFlo Astrios EQ cell sorters (BD Biosciences). Monocytes and granulocytes were cultured in RPMI 1640 medium, supplemented with 10% FCS, 1% P/S, and 2-mercaptoethanol in the presence of recombinant murine granulocyte-macrophage colony-stimulating factor (mGM-CSF) in concentration of 5 or 20 ng/mL for monocytes and granulocytes, respectively, with murine IL-4 (mIL-4;10 ng/mL) added to granulocytes. The inhibitors trametinib (1.642 µM; Selleckhem) and pictilisib (50 nM; Abcam) were added at the specified concentrations and combinations and maintained in the culture for 24 hours.

***In vitro*** **T cell proliferation and activation assays**

When indicated, POM-1 (50 µM), APCP (50 nM; Tocris), anti-PD-1 (2.5 or 5 µg/ml; RPM1-14, ichorbio) or isotype control (2.5 or 5 µg/ml; Rat IgG2A, 1-1, ichorbio), anti-VISTA (50 µg/ml; 13F3 clone; BioXCell) or isotype control (50 µg/ml; PIP clone; BioXCell) were added to the culture and incubated for 4 days.

**Statistics**

Paired two-tailed Student’s t-test was used for normally distributed data, and Mann-Whitney test for non-normally distributed data. Differential gene expression (DGE) analysis was conducted using R packages limma and Voom, with significance defined as fold-change ≥ ±1 and *P* < 0.05. Volcano plots were generated using ggplot2 and ggrepel to visualize log2 fold change and statistical significance (-log10 *P*-value). Heatmaps of molecular expression patterns across clusters were generated using the R package heatmap with Z-score row scaling, or across stem cell subsets using GraphPad Prism 10. Significance levels are indicated as: *p* < 0.05, p < 0.01, *p* < 0.001, p < 0.0001. All statistical analyses were conducted using GraphPad Prism 10.
